# Undiagnosed and diagnosed hypertension in a community setting at Hosanna town: Uncovering the burden

**DOI:** 10.1101/560748

**Authors:** Nebiyu Dereje, Alemu Earsido, Ashenafi Abebe, Layla Temam

**Affiliations:** Department of Public Health, Wachemo University, Hosanna, Ethiopia; Department of Statistics, Wachemo University, Hosanna, Ethiopia; Department of Medical Physiology, Wachemo University, Hosanna, Ethiopia

**Keywords:** Hypertension, Undiagnosed hypertension, Raised blood pressure, Community based, WHO STEPS

## Abstract

**Background:** Hypertension is a leading cause of cardio-vascular diseases and its attributed mortality. No previous study, however, assessed the prevalence and associated factors of hypertension in the study area.

**Methods:** We recruited a representative sample of 627 adult individuals from selected kebeles of Hosanna town. A multi-stage sampling technique was employed in the study. A structured questionnaire using the WHO STEPS approach was employed to conduct a face to face interview and physical measurements. For each participant, we measured blood pressure two times after giving 10 minutes breaks between the measurements and we took the average. Hypertension status was defined as “systolic blood pressure ≥140mmhg and/or diastolic blood pressure ≥90mmhg”. Undiagnosed hypertension was defined as participants who had raised blood pressure on measurement, but not aware of it before. We used Multivariable logistic regression model to determine factors associated with hypertension.

**Results:** The overall prevalence of hypertension was found to be 17.2% (95% CI 14.5 – 19.9), 19.3% among men and 14.2% among women, of which 10.2% were unaware of it before. Hypertension was significantly associated with old age ≥35 years, excess alcohol intake, consumption of saturated oil/fat), consumption of unspecified different types of oil/fat and overweight/obesity.

**Conclusion:** The prevalence of hypertension (both diagnosed and undiagnosed) in the town is unacceptably high. This is also related to modifiable risk factors like excessive alcohol intake, overweight/obesity and consumption of saturated fat/oil. Therefore, designing health information provision systems on the risk factors of hypertension and promotion of good health practices should be considered. Moreover, the health departments should facilitate blood pressure screening programs at community levels to identify and treat undiagnosed hypertension.

## Introduction

Hypertension is a state of high blood pressure and a leading risk factor of cardio-vascular diseases and attributed mortality, globally (1-4). Non-communicable diseases (NCDs) accounted for 72.3% of global deaths in 2016, of which more than 50% of the deaths were attributed by the cardio-vascular problems (1, 2). There is a declining trend of cardio-vascular diseases (CVDs) in developed world due to effective interventions, but, the burden of cardio-vascular diseases is rising in the developing countries, which put the developing countries in the double burden of disease. Ethiopia is not an exception from this rising trend of CVDs (1, 4-6).

In Ethiopia, according to the finding from WHO STEPS survey of 2015, the prevalence of hypertension was found to be 15.8% (7). There are also few studies reported the prevalence of hypertension varying from 8% to 35% (5, 8-16). Moreover, the prevalence of undiagnosed hypertension, those who neither aware of the raised blood pressure nor taking any anti-hypertensive medications. Undiagnosed hypertension may pose serious problem, as it is asymptomatic. Reasons reported for high burden of the hypertensive disorder in Ethiopia were due to life style change, effect of urbanization and globalization (6-11, 17). However, the findings reported were controversial and inconsistent with regard to identifying associated factors of hypertension. The prevalence data is important to understand the magnitude and severity of the problem, identifying high risk groups and measuring effects of interventions. However, the data related to the prevalence of hypertension in the study area is limited and some are hospital (facility based). Therefore, this study describes the prevalence of hypertension and identifies associated factors from the local context, using a community based study design and WHO STEPS approach for surveillance of chronic non-communicable diseases (18).

## Methods and materials

### Study setting, Design and Population

A community based cross sectional **s**tudy was conducted among adult population (>18 years) in Hosanna town, Southern Ethiopia from March 15 to May 20, 2016. Participants were excluded from the study if they were unable to respond for the survey or with mental problem at the time of data collection.

#### Sample size and sampling procedures

Six hundred thirty four (634) samples were estimated by using single population proportion formula; with the assumptions of 95% level of confidence, 5% margin of error, 28.3% prevalence of hypertension in Jigjiga (10) and design effect of 2. Multi-stage sampling technique was employed to recruit samples to be included in the study. In the first step, samples were allocated to the four sub-cities based on proportion to population size. Then 3 kebeles (lowest administrative units in Ethiopia) from each sub-city were selected randomly. Then by consulting the health extension workers and using family folder of the kebele’s population, the final households in the kebeles to be included in the study were identified using simple random sampling. All the eligible individuals in the household were included in the study.

#### Data Collection procedures

Before beginning the data collection, ethical clearance was obtained from Wachemo University ethical review committee and verbal consent of the participants was ensured during the data collection. Face to face interview was conducted by trained nurses by using standardized semi-structured questionnaire which is adapted from WHO STEPS approach instrument. Information on tobacco use, alcohol consumption, fruit and vegetable consumption, physical activity, physical measurement, raised blood pressure, chronic disease history and family health was collected. Formats adapted from WHO STEPS guidelines were also used to measure blood pressure (BP), pulse rate, weight, height, waist and hip circumference.

The research investigators were responsible for the overall management of the project; for development of the final questionnaire, for making the initial contact with and securing participation of the kebeles included, for identifying survey administrators and to train and assign them to the selected kebeles. The Data collectors were ten trained nurses who were supervised by two recruited Bsc. Public health professionals who were working in the study area at the time of data collection.

Face to face interview was conducted at home level after the interviewers explained the purpose of the study and obtained the participant’s informed consent to participate in the study. Eligible respondents were declared unavailable if they were not found on three separate visits. After completion of face to face interview, all respondents were given appointment for physical measurements and it was taken in the outreach sites inside the kebele, which is conducive for the community to be involved in the study. All study instruments were translated into local languages (Amharic) by native speakers and then back translated to English by other persons who understand the languages, to see the consistency. Pretest was conducted in 5% of the samples to see the completeness, consistency, and applicability of the instruments and was ratified accordingly. Instruments used for measuring the physical dimensions like weight scale, and height measuring board were calibrated in a daily basis and checked after every measurement. Daily supervision was made in the field during data collection by field supervisors and investigators. Data collectors have checked for data completeness and consistency before leaving each house. Field supervisors also have checked the completeness and consistency of the data on daily basis and they returned to interviewers if the data were incomplete and inconsistent.

#### Measurement of variables

**Height** is measurement of the body physical dimension between the top of the head and the bottom of the feet. The height was read in centimeters to the nearest 0.1cm and recorded. **Weight** was measured using digital weighing scale and recorded in kilograms to the nearest 0.1kg. **Hip to waist circumference ratio measurement** is the circumference of the waist measured mid-way between the lowest rib cage and anterior superior iliac spine divided by the circumference of the hip measured at the level of the greater trochantor of the fumer (both are measured to the nearest 0.1 cm). **Blood Pressure** – the measurements was taken using digital blood pressure measuring apparatus. Two measurements were taken for analysis purposes; recording the mean of the first and second readings. The right arm was used for this measurement. The displayed reading of the systolic and diastolic blood pressure was recorded. Participants have taken rest for ten minutes between each reading. Hypertensive status of the participants was defined as systolic blood pressure ≥90mmhg and/or diastolic blood pressure ≥140mmhg. Undiagnostic hypertension was defined as participants whose systolic blood pressure ≥90mmhg and/or diastolic blood pressure ≥140mmhg and unaware of it before. **Tobacco use** was defined as using tobacco products within the last one month preceding the survey. **Harmful use of Alcohol** was defined as consumption of ≥ 60 gm of pure alcohol (6 drinks) on average per day for men and ≥ 40 g (4 drinks) for women. **Physical activity** was measured by estimating the mean physical activity (PA), which was calculated as metabolic equivalent in minutes per week and respondents’ physical activity. Then it was categorized as low, moderate and high. The domains investigated activity at work, in travel, sports, fitness and recreational/leisure. **Unhealthy diet** is consumption of unhealthy diet was measured using the fruit and vegetables consumption level, eating outside the home and type of oil/fat most used for meal preparation.

#### Data management and analysis

Data were checked, cleaned, and entered in to Epi data 3.1. Version software, then imported to SPSS version 20 software for analysis. Incomplete and inconsistent data were excluded from the analysis. Descriptive statistics were used to describe the sample. The prevalence of hypertension was described using the proportion and 95% confidence interval. Associations between independent variables and dependent variables were analyzed first using bivariate analysis to identify factors eligible for multivariable analysis. Those variables with p-value < 0.25 in the bivariate analysis were included in the multi-variable analysis. The magnitude of the association between the independent and dependent variables was measured using odds ratios (OR) and 95% confidence interval (CI) and P values below 0.05 was considered statistically significance.

## Results

### Socio-demographic characteristics of the respondents

A total of 627 respondent’s data were analyzed, with the response rate of 98.9%. Among them 58.5% were males and 41.5% were females and majority of them were in the age category of 25- 34 years (32.4%). The mean and median age of participants was 36 and 34 years respectively. Majority of the respondents are married (65.6%) and self-employed (31.6%). Of all the respondents approached, 95% indicated that they had formal education and able to read and write, but the remaining 5% of the respondents can’t read and write (**Table 1** – Socio-demographic characteristics of the respondents in Hosanna town, March 15 to May 20, 2016 (n=627).

**Table 1.** Socio-demographic characteristics of the respondents in Hosanna town, March 15 to May 20, 2016 (n=627)

| Variable | Response | Frequency (percent) |
| --- | --- | --- |
| Sex | Male | 367(58.5%) |
|  | Female | 260(41.5%) |
| Age | <25 years | 112(17.9%) |
|  | 25-34 years | 203(32.4%) |
|  | 35-44 years | 155(24.7%) |
|  | 45-54 years | 105(16.7%) |
|  | 55-64 years | 40(6.4%) |
|  | ≥65 years | 12(1.9%) |
| Ethnicity | Hadiya | 397(63.3%) |
|  | Kembata | 121(19.3%) |
|  | Gurage | 32(5.1%) |
|  | Silte | 20(3.2%) |
|  | Amhara | 36(5.7%) |
|  | Others | 21(3.3%) |
| **Marital status** | Single | 216(34.4%) |
|  | Married | 411(65.6%) |
| **Educational status** | Illiterate | 32(5.1%) |
|  | Literate | 595(94.9%) |
| **Occupation** | Government employed | 181(28.9%) |
|  | NGO employed | 38(6.1%) |
|  | Self employed | 198 (31.6%) |
|  | Student | 59(9.5%) |
|  | Housewife | 87(13.9%) |
|  | Retired | 21(3.3%) |
|  | Unemployed | 43(5.8%) |

### Prevalence of Hypertension

Blood pressure of the respondents was taken two times per individual and the average was recorded. The mean systolic blood pressure of the respondents was 123.4 mmhg and the mean diastolic blood pressure was 75.3 mmhg. With regard to their hypertensive status (which is defined as systolic blood pressure of greater than or equal to 140 mmhg and/ or diastolic blood pressure of greater than or equal 90 mmhg), 17.2% were hypertensive (95%CI 14.5 – 19.9), which is 19.3% among men and 14.2% among women. Whereas, about 10.2% of those with raised blood pressure were not aware of it before. Moreover the study has revealed that the occurrence of hypertension is increasing in a linear manner with increase in the age of the respondents (**Fig 1**).

**Figure 1.**
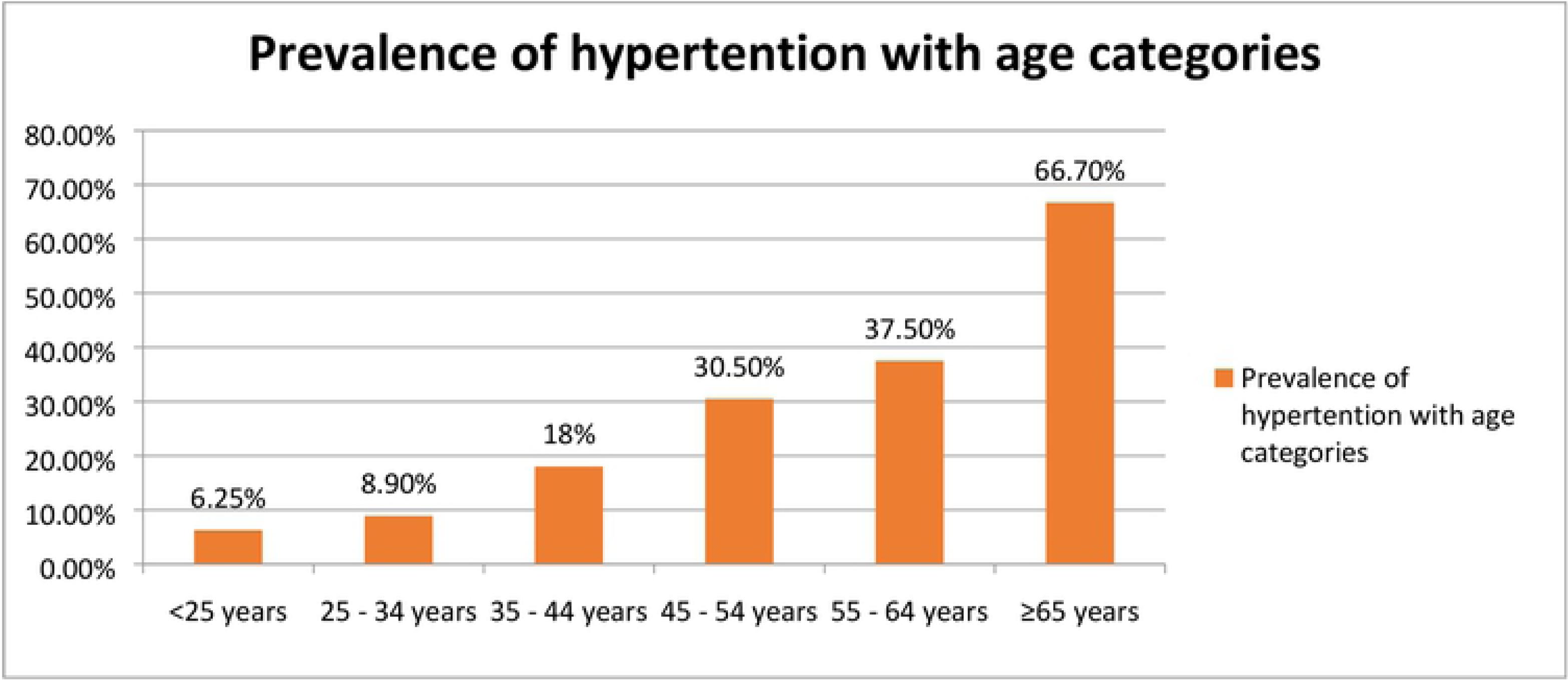
Prevalance of hypertension in relation to the age category of respondents in Hosanna town, May 2016

### Behavioral and dietary risk factors of hypertension

Overall, 29 (4.6%) of the survey population indicated they are current smoker of tobacco products. Among them, 11 (38%) were hypertensive (p – value <0.005). The proportion of current alcohol drinkers (alcohol consumption in the past 30 days) was 15.9%, which was 21% among men and 8.8% among the women. Majority of the current alcohol drinkers were in the age category of 35-44 years. Among the current alcohol drinkers, 34 (34%) of them were hypertensive (P-value <0.0001). In the last week preceding the survey, about 86 (13.7%) of respondents ate no fruit and/or vegetables on an average day in the last week preceding the survey. Among them, 21 (24.4%) were hypertensive (P-value <0.05). High proportion of the respondents 317 (50.6%) used saturated fats (oils) for meal preparation (margarine, lard and butter), whereas about 232 (37%) used unspecified different types of fats (oils) for meal preparation. However, only 78 (12.4%) used vegetable oils for meal preparation. Moreover, about 30% of the participants had more than two times meal outside of their home in the last week preceding the survey. Among the respondents who used saturated fats and unspecified different types of fats for meal preparation, 56 (17.7%) and 49 (21.1%) were hypertensive respectively (P – value <0.001). With regard to the participants’ physical activity, only 6.1% of the respondent’s work involves vigorous intensity activity and about a quarter (28.4%) of the respondents was engaged in a work requiring moderate intensity activity, whereas the remaining 65.5% of the respondent’s work involves low intensity physical activity. Among the vigorously working group, they work on average of 4.73 days for a mean spent time of 4.78 hours. Likewise among the moderately working group, they work on average of 4.38 days for a mean spent time of 3.64 hours. The study also indicated that about a quarter 24.8%, (57% among males and 43% among females) of the respondents involved in vigorous intensity sports, fitness, or recreational activities which causes large increase in breathing and heart rate, where they spend an average 2.06 days per week and 50.79 minutes per the activity. On the other hand, the respondents spend on average of 5.63 hours per day by sitting. About 60 (9.5%) of the respondents spend more than 8 hours per day in a sitting (sedentary life style). Among them, 19 (31.7%) were hypertensive (P value < 0.001).

Mean height of the respondents were 169 cm, which is 171 cm among men and 165 cm among women. The mean weight of the respondents was 66.98 Kg, which is 68.4 Kg among men and 64.6 Kg among women. The mean BMI of the respondents was 23.63 kg/m^2^, which is 23.46 kg/m^2^ among men and 23.87 kg/m^2^among women. Majority of the respondents were with normal weight (72.8%) and about 23.7% of them were overweight and only 1.3% were obese on the BMI category. about one third of overweight and a quarter of obese participants were hypertensive (P-value <0.001). Waist circumference is a measure of central obesity, which is the type of obesity that predisposes to the chronic non-communicable diseases. A waist circumference that is greater than 94cm (37inches) in males and 80cm (31.5 inches) in females defines central obesity. In this study, the mean waist circumference in men was 91.3 cm, and for women it was 83.3 cm, which is above of 80cm. The mean waist circumference for women was higher in the age group 55 – 64 years (mean 88.7 cm). The mean hip circumference was found to be 102.2 cm in men and 109.4 cm in women. Another measure of central obesity is the waist/hip ratio. In men, one is said to be obese if it is above 1 and in women if it is above 0.8cm. On average the waist /hip ratio was found to be 0.89 and 0.76 cm among men and women respectively.

In the bivariate analysis, variables which were significantly associated with hypertension were: age ≥35 years, current smoker, who are currently drinking more than 5 bottles of alcohol in the week, the type of oil used, spending more than 8 hours per day in a sitting position and abnormal BMI. However, being male sex and not consuming vegetables and/or fruit in the last week didn’t shown significant association with hypertension, but they were included in the multi-variable analysis, since they had P-value below 0.25.

In multi-variable analysis considering those variables having P-value < 0.25 in bivariate analysis, hypertension was significantly associated with age ≥35 years, which are currently drinking more than 5 bottles of alcohol in the week, the type of oil used, and abnormal BMI (**Table 2** – Factors associated with hypertension in Hosanna town,). Older respondents (age > 35 years) were 4 times more likely to be hypertensive than the younger ones (AOR = 3.97, 95% CI 1.45 – 10.83). Those individuals who drink more than five alcoholic drinks (> 60gm)/day were 3 times more likely to be hypertensive as compared to those who drink less than five alcoholic drinks per day (AOR = 2.9, 95% CI 1.1 – 7.6). Individuals who used unsaturated oil (AOR = 6.5) and unspecified different types of oils (AOR = 8.2) for cooking meals were more likely to be hypertensive than those who used vegetable oils for cooking meals. Overweight/obese individuals were 3 times more likely to be hypertensive as compared to the normal weighted individuals (AOR = 2.7, 95%CI 1.7 – 4.3).

**Table 2.**
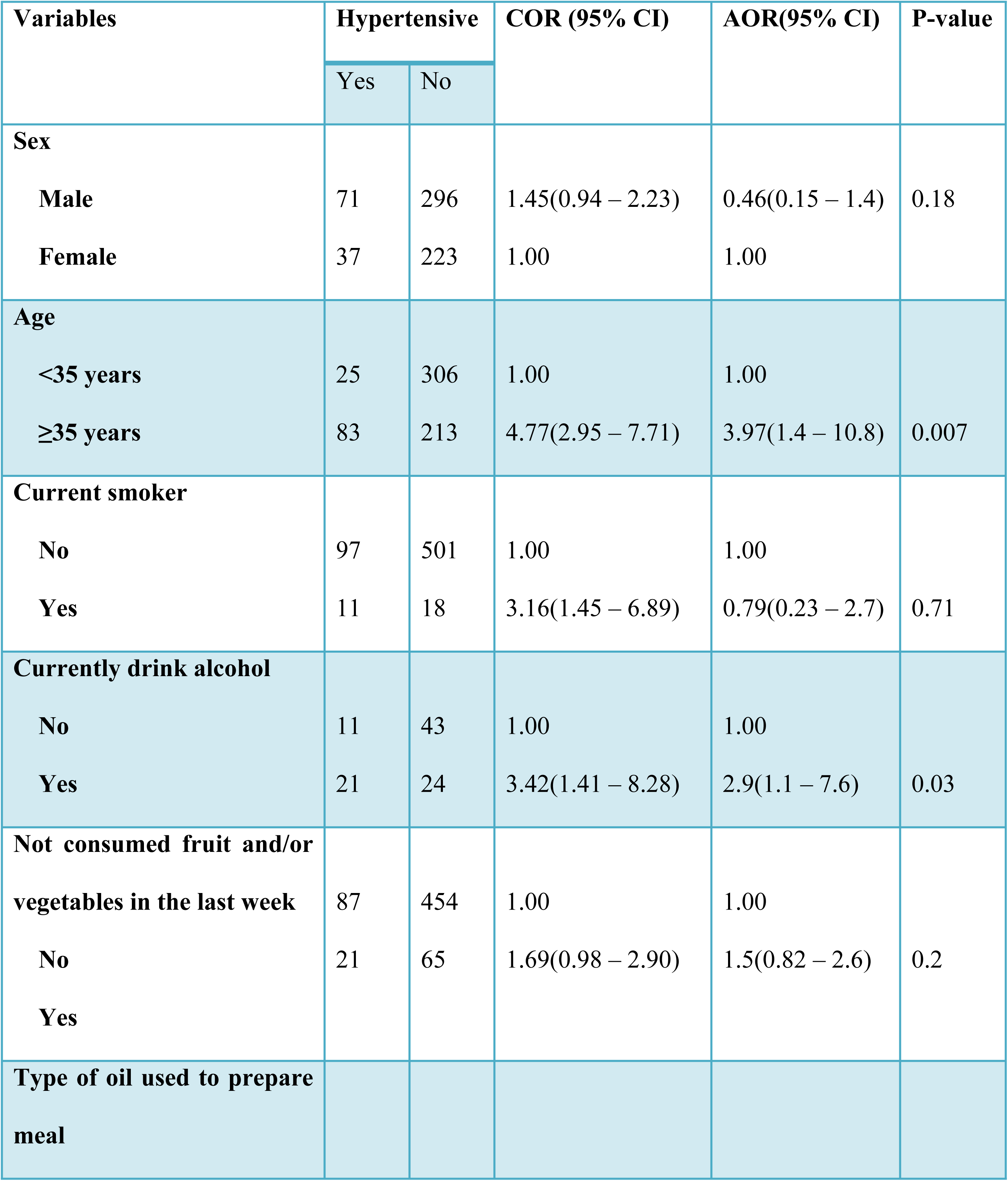
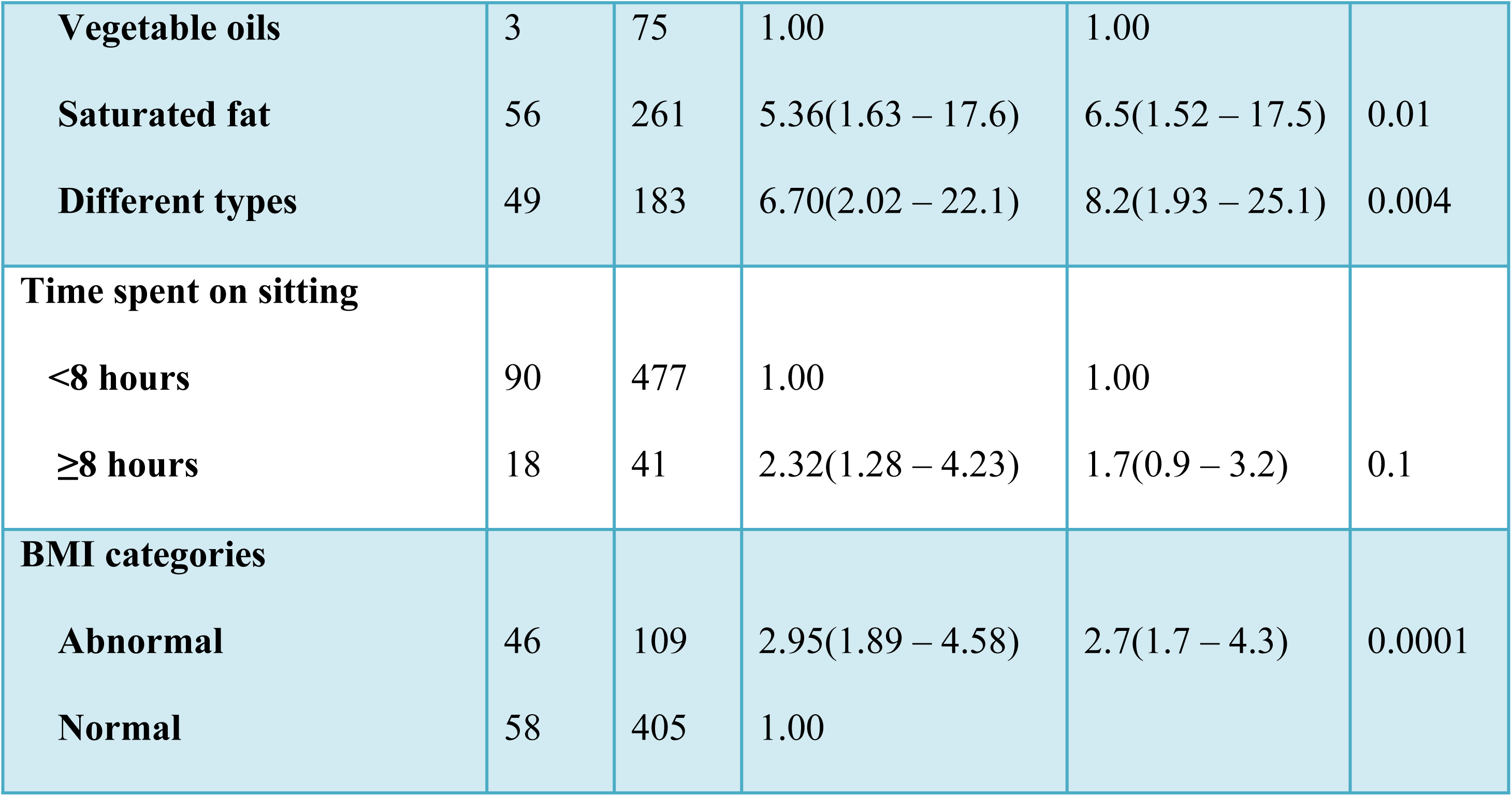
Factors associated with hypertension in Hosanna town, March 15 to May 20, 2016 (n=627)

## Discussion

The burden of hypertension and its associated cardiovascular problems are increasing in developing countries like Ethiopia (1, 3, 4). The present study found out the overall prevalence of hypertension to be 17.4%. More than half of the respondents were not aware of their hypertensive status (10.2%). Hypertension (high blood pressure) is asymptomatic until complications arise and may cause serious health problems which include sudden death due to cardiac problems, if left unrecognized and untreated. High proportion of undiagnosed (unaware) hypertension calls for different public health interventions like providing hypertension screening programs at the community level (19-21).

The finding of the present study is consistent with the findings on the prevalence of hypertension in Hawassa (19.7%) (8) Mekelle (19.1%) (5) and the national prevalence (15.8%) (7). However, it is lower than the findings of the studies in Addis Ababa (32.3%) (15), Jigjiga (28.3%) (10), Durame (22.4%) (12), Bahir dar (23.5%) (22) and Gonder (28.3%) (13) and higher than the findings of the studies in Jimma (13.2%) (11) and Sidama zone (9.9%) (9). The discrepancy in the prevalence of hypertension might be attributable to the settings of the studies, the age group of the participants included in the studies and the life style of the population in the study areas. This study revealed that hypertension is associated with old age, excessive alcohol drinking, utilization of saturated fats/oils or unspecified different types of fats/oils and abnormal BMI. It has been mentioned by other reviewed studies that an increment in age has the effect on the blood pressure, which might be due to changes that happen in the walls of blood vessels as age increases (5, 10, 11). Similarly, consumption of excessive saturated fat/oil is also a risk factor for hypertension. This is because; the body will convert saturated fats into cholesterol, which in turn will narrow the arteries and raises resistance in the blood vessels, resulting in high blood pressure [(23)]. Studies also identified that abnormal BMI (obesity) is associated with hypertension (5, 10, 23).

The present study has employed a community based design, which allows generalization to the population of the town. Moreover, the study included participants’ interview and physical measurements using standard procedures, which allowed us to triangulate the study findings from different sources. However, this study has limitations since it has employed cross sectional study design and some of the variables were taken for a study period only. For instance, nutrition related questions were assessed for one week preceding the survey and might not represent the usual pattern of life styles and also prone to recall bias.

## Conclusion

In sum, the prevalence of both diagnostic and undiagnosed hypertension in the town is unacceptably high. This is also related to modifiable risk factors like excessive alcohol drinking, saturated fat/oil consumptions and physical inactivity. Therefore, due consideration should be given for prevention and control of hypertension by designing health information provision systems on the risk factors of hypertension and promotion of good health practices. Moreover, the health departments should facilitate blood pressure screening programs at community levels to identify and treat undiagnosed hypertension.

## Acknowledgement

We would like to thank Wachemo University for the funding of this study. We would also like to express the study participants and data collectors for their contributions for the success of this study.

## Source of Funding

Funding of this study was obtained from Wachemo University.

